# Single variant bottleneck in the early dynamics of *H. influenzae* bacteremia in neonatal rats questions the theory of independent action

**DOI:** 10.1101/075531

**Authors:** Xinxian Shao, Bruce Levin, Ilya Nemenman

**Affiliations:** Department of Physics, Emory University, Atlanta, GA 30322, USA; Department of Biology, Emory University, Atlanta, GA 30322, USA

**Keywords:** bacterial infection, single–variant bottleneck, phenotypic switching, stochastic processes

## Abstract

There is an abundance of information about the genetic basis, physiological and molecular mechanisms of bacterial pathogenesis. In contrast, relatively little is known about population dynamic processes, by which bacteria colonize hosts and invade tissues and cells and thereby cause disease. In an article published in 1978, Moxon and Murphy presented evidence that, when inoculated intranasally with a mixture streptomycin sensitive and resistant (Sm^*S*^ and Sm^*R*^) and otherwise isogenic stains of *Haemophilus influenzae* type b (*Hib*), neonatal rats develop a bacteremic infection that often is dominated by only one strain, Sm^*S*^ or Sm^*R*^. After rulling out other possibilities through years of related experiments, the field seems to have settled on a plausible explanation for this phenomenon: the first bacterium to invade the host activates the host immune response that ‘shuts the door’ on the second invading strain. To explore this hypothesis in a necessarily quantitative way, we modeled this process with a set of mixed stochastic and deterministic differential equations. Our analysis of the properties of this model with realistic parameters suggests that this hypothesis cannot explain the experimental results of Moxon and Murphy, and in particular the observed relationship between the frequency of different types of blood infections (bacteremias) and the inoculum size. We propose modifications to the model that come closer to explaining these data. However, the modified and better fitting model contradicts the common theory of independent action of individual bacteria in establishing infections. We discuss the implications of these results.

## 1. Introduction

Before the *Hib* (*Haemophilus influenzae* type b) conjugate vaccine was developed and taken into routine use in the U. S., *H. influenzae* was the leading cause of bacterial meningitis in children under the age of five [1]. At the same time, bacterial meningitis had high mortality and serious sequela, including deafness, blindness and mental retardation. Even today, at least in part because of the lack of vaccines, in the developing world the mortality rate from *H. influenzae* infections is substantial, with case mortalities approaching 14.3% [1].

In 1974 Richard Moxon and colleagues developed a neonatal rat model to study the pathogenesis of *Haemophilus influenzae* [2]. They presented evidence that *Hib* infection could be divided into three elements: nasopharyngeal colonization, bacteremia, and central nervous system (CNS) invasion. In 1978, by intranasally inoculating the neonatal rats with mixtures of otherwise isogenic streptomycin sensitive and resistant strains of *H. influenzae*, Sm^*S*^ and Sm^*R*^, and tracking the development of bacteremia and meningitis, Moxon and Murphy found that, five minutes after inoculation, *both variants* were found in the blood. In contrast at 54 hours, nearly 70% of rats had *pure* Sm^*S*^ or Sm^*R*^ in the blood (Figure 1) and cerebrospinal fluid, and the cultures had Sm^*S*^ and Sm^*R*^ isolated in nearly equal frequency [3]. Their observation that the primary infections (nasal colonization and early blood flora for *Hib*) are diverse, while mature infections (blood after *>* 10 hrs for *Hib*) were monoclonal is known as the *single-variant bottleneck*. The bottleneck is not unique to *Hib*. In fact, it was discovered first by Meynell in *Salmonella typhimurium* infections in mice [4]. And similar observations have been reported in experimental studies of other host-bacteria [5, 6, 7] and host-virus [8, 9, 10, 11, 12, 13, 14] interactions.

**Figure 1.**
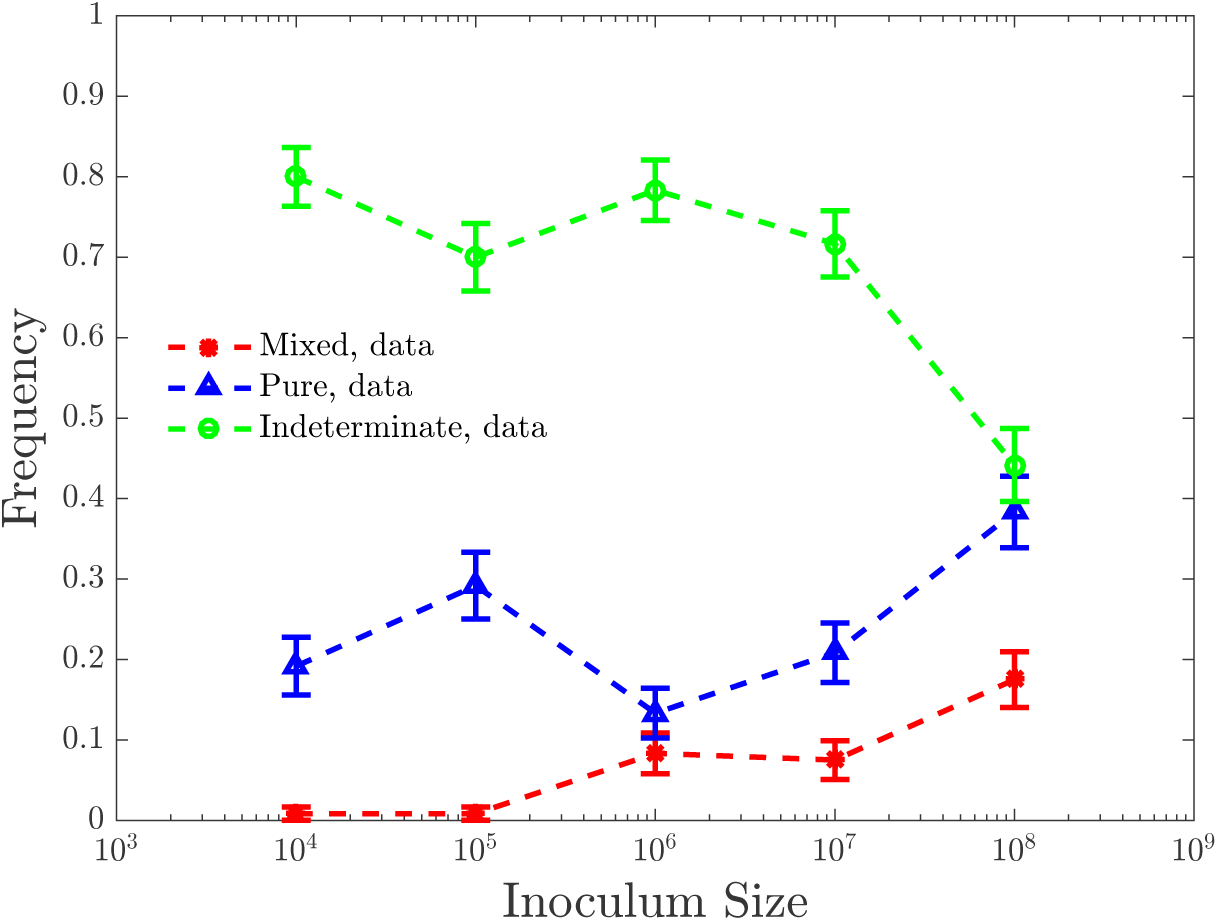
Moxon and Murphy’s experimental data. Replotted from [3]. 120 neonatal rats were infected at each inoculum size, which ranged from 10^4^ to 10^8^ bacteria, equally mixed from streptomycine susceptible strain, Sm^*S*^, and streptomycine resistant strain, Sm^*R*^. Blood of the rats was then harvested and cultured. All cultures that produced both Sm^*S*^ and Sm^*R*^ colonies were called *mixed* infections. All cultures that produced 8 or more colonies of one strain and none of the other were called *pure* infections (there was no statistically significant difference in the abundance of pure Sm^*S*^ or pure Sm^*R*^ cultures). Cultures that produced no colonieshown on the plots, or produced colonies of one strain only, but fewer than 8 of those, were called indeterminate. Most infections ended up as pure (single-variant) infections 54 hours post-inoculum. Samples taken within 5 and 30 minutes post-inoculum were mixed (data not show on the plot), see Ref. [3]). Error bars denote the usual square-root counting errors. Over four orders of magnitude in the inoculum size, preponderance of indeterminate infections declined somewhat, from ∼80% to ∼40%. Over the same range of the inoculum, mixed infections increased from <1% to ∼20%.

Models of the infection process must be able to explain the single-variant bottleneck, and also the fact that, although there are high frequencies of people colonized with virulent strains of *H. influenzae* as carriers, only minority of hosts (e. g., children under 5 [3, 1]) manifest invasive infections even for the most virulent strains of these bacteria. Three classes of possible explanations have been proposed: stochasticity resulting from *independent action* of bacteria [4, 15, 16, 17, 18], variations in host susceptibility, and the emergence of bacterial mutants (known as within-host evolution) [19, 20, 21]. The original theory of independent action was proposed by Druett in 1952 [22]. It assumed that each individual bacterial cell has an independent probability to colonize the host, and that an infection can start from a single random bacterium, hence explaining the bottleneck. Meynell and Stocker later provided experimental evidence consistent with the theory of independent action [4] and inconsistent with an alternative: synergistic or cooperative action. However, in its simple form, independent action could not explain why a variant present in the blood five minutes post-inoculation can no longer be detected a few hours later. Host susceptibility could only explain the high infection rate in young children over adults, but not the experimental observation of the presence of both variants in the bloodstream only very early post-inoculation stage [3]. Finally, to test the within-host evolution hypothesis, Margolis and Levin performed additional *H. influenzae* experiments with neonatal rats. They compared the invasiveness and the re-colonization potential of the variant surviving in the bloodstream and the remaining variant staying in nasopharynx [21]. In five out of six clones examined, they observed no difference in the ability of bacteria isolated from the blood to outcompete those remaining in the nasopharynx during re-infection. This result suggested that within-host evolution is not the dominant explanation for the Moxon-Murphy observations.

During the last two decades it has become clear that bacteria can switch phenotypic states without any changes in their genomes [15, 17, 23, 24, 25, 26, 27]. This provides an alternative hypothesis to explain the Moxon-Murphy results. First, a single bacterium of one strain can randomly switch into a faster growing phenotype or a more invasive state. Then the host immune system, activated by the infection emerging from the switched bacterium, acts against both strains and clears the clone that has not phenotypically switched and thus grows slowly. In other words, the first successfully switching cell in one strain will interact with the host immune response to ‘shut the door’ on the other strain and thereby explain the single-variant bottleneck and the subsequent failure to isolate genetically more invasive clones from the blood [21].

This stochastic switching mechanism together with the immune response has been mentioned frequently as a possible explanation for the bottleneck phenomenon in various presentations and discussions. Nevertheless, we have been unable to find a detailed theoretical or experimental analysis of this process in the literature. We call this phenotypic switching mechanism followed by immune-facilitated clearance of the competing strain the *colloquial hypothesis*. Our goal here is to analyze the colloquial hypothesis quantitatively and to explore the condition under which it can or cannot rescue the theory of independent action as the explanation of the single-variant bottleneck in early bacteremia.

In the following sections, we will develop a mathematical model of the colloquial hypothesis applied to the early stages of *Hib* bacteremia inoculated with two variants of equally invasive bacteria. We will show that the model, as well as its simple extensions, cannot provide a quantitative explanation for the experimental data [3, 20, 21, 28], and namely the observed weak dependence of the rate of different types of infections on the inoculum size. We argue that, to provide even a semi-quantitative fit to the data, we must assume that various rate parameters describing the infection scale *sublinearly* with the inoculum size, so that the probability per bacterium to start an infection decreases when other bacteria are present. This means further evidence for abandoning the theory of independent action.

## 2. Hypothesis and Model

Inspired by demonstration of phenotypic switching in bacteria [17, 24, 25, 26, 27, 29, 30, 31], we propose that each individual bacterial cell of both variants A and B has two phenotypes relevant for the early infection. The first is the “crossing” phenotype (C), which allows bacteria to cross the physical barrier between the nasopharynx and blood, but does not exhibit strong growth in the bloodstream [32, 33, 34]. The second is the “growing” phenotype (G), with cells that grow fast in the bloodstream, but cannot cross into the bloodstream. After a bacterium crosses into the bloodstream, it can switch to the G state, but the switching C→G is stochastic and rare. In this work, we are not concerned with the mechanisms underlying the existence of these two states and of switching between them, but only focus on consequences of the switching.

Once bacterial cells enter the bloodstream, immune response is activated. To model the immune response in the early stages of bacteremia, we assume that neonatal rats only have innate immunity, which is non-specific and responds as soon as the bacterial cells emerge in the blood [1, 3, 19, 35]. In other words, there is no clonal expansion, and instead there is a finite reservoir of immune cells that can be recruited to the infection site linearly until the reservoir is depleted [35].

These assumptions are represented in the following ordinary differential equations (ODEs) describing the growth of bacterial cells of variant A and the immune cell recruitment:

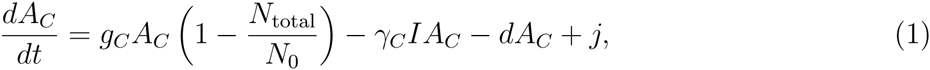

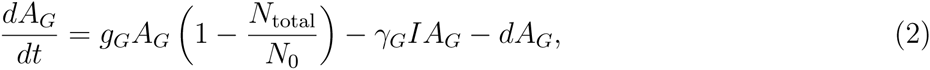

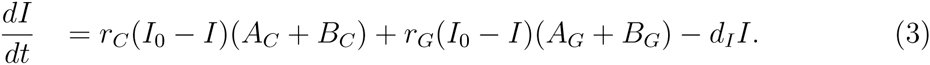

The growth of the bacterial strain B is described by equations similar to Eqs. (1, 2), with letters A replaced by B.

In the equations describing bacterial population dynamics, Eqs. (1, 2), *g*_*C*_ and *g*_*G*_ are the growth rates of the crossing and the growing phenotypes, respectively (same for variants A and B since both variants are equally virulent [3, 21]). For simplicity, in what follows we set *g*_*C*_ = 0. *N*_total_ is the total number of bacteria in the blood, *N*_total_ ≡ *A*_*C*_ + *A*_*G*_ + *B*_*C*_ + *B*_*G*_. *N*_0_ is the carrying capacity, the maximum of the bacterial population in the bloodstream, so that *N*_0_ ≥ *N*_total_. *γ*_*C*_ and *γ*_*G*_ are the bacterial death rates due to the elimination by the immune cells, and *d* is the natural cell death. Finally, *j* is the flux of the crossing phenotype cells from the nasopharynx to the bloodstream per unit time. To satisfy the hypothesis of independent action, it is assumed to be linearly proportional to the inoculum size *S*, so that *j* = *α*_*j*_*S*, where *α*_*j*_ is some constant.

Equation (3) describes the immune cells recruitment. Here *r*_*C*_ and *r*_*G*_ are the recruitment rates due to effects of the bacterial phenotypes [35]. *I*_0_ is the total number of available innate immune cells in the host. *d*_*I*_ is the death rate, or deactivation rate of immune cells. Parameters in Eq. (3) are determined up to a scale. Thus we set *I*_0_ = 1, which redefines the scale of *I* and also renormalizes *γ*_*C*_ and *γ*_*G*_ in Eqs. (1, 2). To simplify the model, we set *γ*_*C*_ = *γ*_*G*_ = *γ*, and *r*_*C*_ = *r*_*G*_ = *r*.

To finish specifying the model, we assume for now that the switching from C to G is a single step stochastic transition at a low per-bacterium rate *ρ*. Since the switching is single-step, the waiting time to the switch is exponentially distributed for each cell. Further, if the independent action hypothesis is valid, then for *A*_*C*_ bacteria in the crossing phenotype, the probability of having *k* individuals of type A switching to the growing state per time ∆*t* is given by the Poisson distribution:

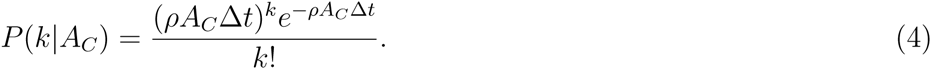

A similar distribution determines the switching probability for the B strain. We do not consider switching back from the G to the C state.

We simulate the model using the Euler method to solve its ODEs, Eqs. (1, 2, 3) and their equivalents for the B strain. Further, at every time step, we generate a random number of switching individuals using Eq. (4) for the A and the B strain. If the number of cells in any strain / phenotypic state combination falls below one, we set it to zero to account for the discreteness of the bacteria. While this combined stochastic-deterministic simulation scheme is certainly not the most accurate, we feel that it offers the precision necessary for our analysis. It certainly is capable of discovering the salient qualitative features of the models that we investigate. Further, it is much faster that fully stochastic simulations schemes, which is important since the model must be solved repeatedly in optimization steps. Finally, statistical properties of *Hib* bacterial division are not well understood; thus building a fully stochastic model of the system, including modeling the divisions as first order, memoryless reactions, would be not any more accurate than neglecting the stochasticity altogether.

## 3. Results

### 3.1. The colloquial model

We illustrate a possible dynamics of the colloquial model in Fig. 2 for the first 60 hours post-inoculum in an individual rat with a certain set of model parameters. In this case, a cell of strain B switched to the growing phenotype first at *t* ≈ 11 hrs. Then the rapid growth of *B*_*G*_ accelerated the recruitment of immune cells. In their turn, the immune cells nearly wiped the population of the non-switched strain A, transforming the infection into the pure B infection about 30 h post-inoculation. Therefore, even though cells act independently, they interact through the immune response, and the first variant to have a switcher wins the competition. This example illustrates that the colloquial hypothesis may have a potential to explain the single-variant bottleneck in the early stages of bacteremia.

**Figure 2.**
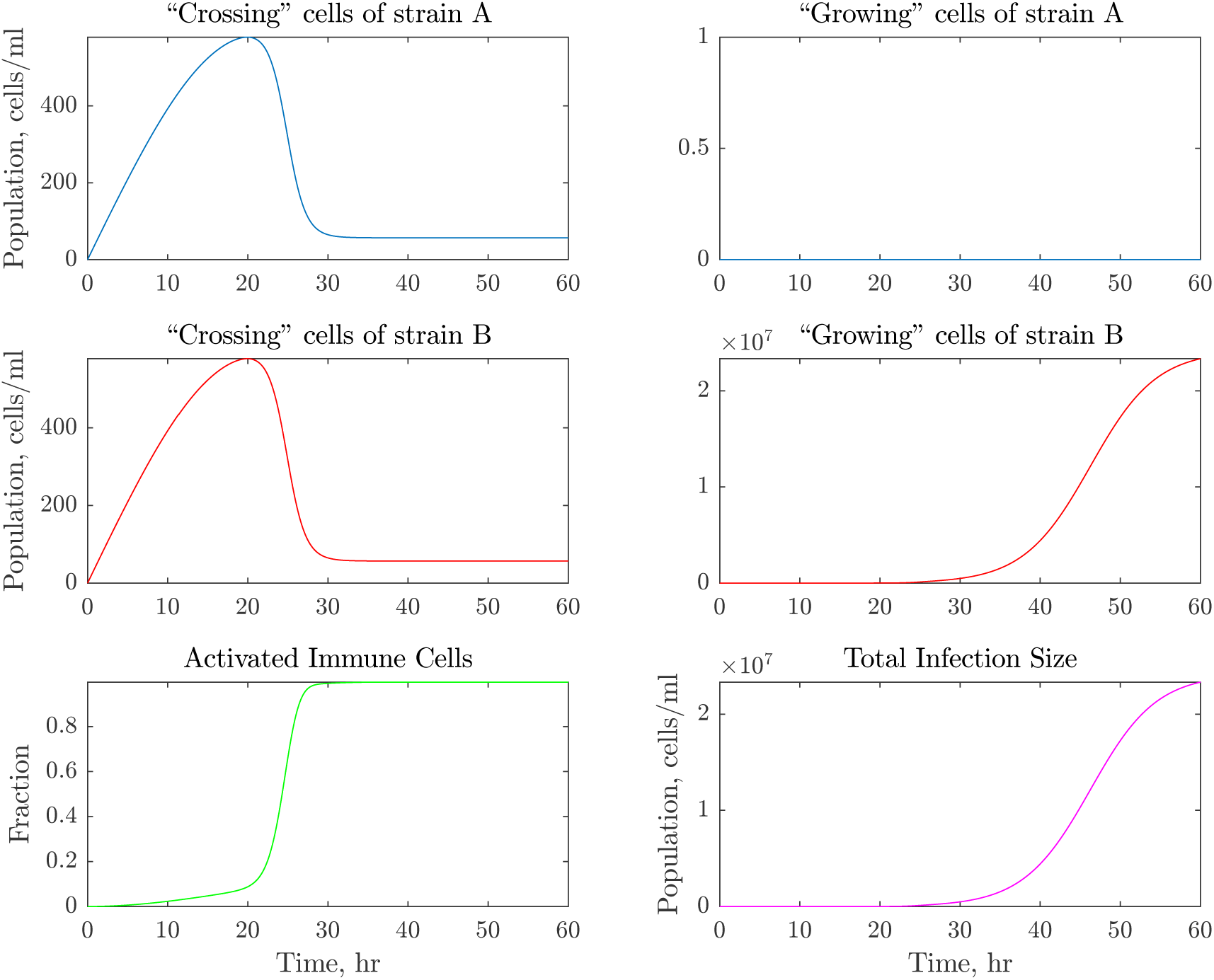
Simulation of early bacteremia resulting in a pure infection in an individual rat. Simulations were done with the following parameters: inoculum size *S* = 10^5^; switching rate *ρ* = 4.5 · 10^−5^ h^−1^; growth rates *g*_*C*_ = 0, *g*_*G*_ = 1 h^−1^; immune recruitment rates *r* = 6.1 · 10^−6^ h^−1^; rate at which immune cells kill bacteria *γ* = 0.75 h^−1^; carrying capacity of the blood *N*_0_ = 10^8^ cells; flux from the nasopharynx to blood *α*_*j*_ = 4.3 · 10^−4^ h^−1^; natural death rate of bacteria and immune cells *d* = 0.01 h^−1^ and *d*_*I*_ = 0.02 h^−1^. In this realization, variant *B* has the first switch from *C* to *G* at about 11 hours and establishes bacteremia. The panels show the population size of the crossing and the growing phenotypes of A and B strains, the fraction of the immune response activated, and the total infection size. Notice that the vertical axes in different panels have different scalings.

To test the suitability of the colloquial model quantitatively, we calculate and maximize its likelihood given the observed experimental data. As in the Moxon and Murphy experiment, we assume the multinomial structure of the data with three possible outcomes: pure infection, mixed infection, and indeterminate infection. Recall that Moxon and Murphy plated blood samples from their rats and counted the number of colonies of each strain that grew as a result. They defined any culture with colonies of both strains (even if one of the strains had as few as one colony) as a mixed infection. A pure infection was defined more stringently, so that there had to be at least eight colonies of one strain and none of the other to qualify. All other cases were deemed indeterminate. In our simulations, accounting for dilution at plating, we estimate that a mixed infection would require both bacterial strains present at a level of 100 cells/ml of bacteria or more, and a pure infection would require at least 800 cells/ml of bacteria of one type and less than 100 cells/ml of the other. To calculate the likelihood of the data given a set of parameters, we simulate infections using our mixed stochastic-deterministic simulations. We repeat this 200 times to estimate the multinomial probabilities. We then write down the multinomial likelihood of the experimental data given the frequencies defined by the numerical simulations. Finally, we optimize the model over the parameters using patternsearch from Matlab with GPS Positive basis Np1 as the poll method. This routine allows optimization of stochastic functions. The optimization is performed at least three times from different initial conditions, and we report the best fit model as the one maximizing the likelihood over all such optimization runs.

The experimental data that we fit contains five different inoculum sizes, and three possible outcomes at 54 hrs post inoculation (for a total of 10 independent data points since the frequencies at each inoculum sum to one). In addition, the experimental data contains measurements a few minutes after inoculation for each inoculum size (10 more independent data points), at which point *every* infection was mixed. Note that these data are not time series—every rat could be analyzed only once—so that the data at different time points are independently multinomially distributed.

These 20 data points must be explained by 8 independent parameters: *α*_*j*_, *g*_*G*_, *d*,*N*_0_, *r*, *γ*, *d*_*I*_, and *ρ*. This may sound like an easy fitting problem. However, it turns out that the requirement of having all mixed infections soon after inoculation, and a lot of pure infections later on is not easy to satisfy. Thus we do not perform formal analysis of the quality of fit / overfitting in this and the other models we try: the difficulty to fit the data makes most models obviously poor, and differences among the quality of fits of various models are clear without formal analyses.

The optimization is further constrained since biologically realistic limits exist on the model parameters. First, all of the parameters are positive. Further, an upper limit on *N*_0_ is about ∼ 2 · 10^9^ cells/ml [36], while its lower limit is determined by the fact that Moxon and Murphy observed ∼ 1 × 10^4^ or more cells/ml in the bloodstream of neonatal rats with severe infections [3]. The growth rate of *Hib* in synthetic blood culture was studied in [36], which provides the initial guess and upper limit of *g*_*G*_ between 0.4 to 1.2 per hour. We could not find any data in the literature regarding the parameters of the innate immune response to *Hib*. However, some data is available for *Listeria* infection [37, 38], which allowed us to choose initial conditions of the immune response for the optimization: *r* ∼ 1 · 10^−6^ h^−1^ cells^−1^, *γ* ∼ 0.1 h^−1^, *d*_*I*_ ∼ 0.02 h^−1^.

Two of the best quantitative fits of the colloquial model are shown in Fig. 3. Some of the parameters of these fits are physiologically unrealistic, but even this does not help: none of the fits are good. The main difficulty seems to be that keeping the fraction of pure / indeterminate infections nearly constant over four orders of magnitude of the inoculum size, *S*, especially following a mixed infection soon after inoculation, is impossible within this independent action model. Indeed, the fit in the left panel keeps pure infections at nearly zero frequency in order to have few mixed infections. Similarly, in the right panel, which does a better job in fitting the frequency of pure infections, the mixed infections rate spikes to 100% at high *S*. Note parenthetically that the non-monotonicity of the mixed infection line in this figure is because of the linearly increasing flux *j* interacting with the immune system. When *S* = 10^4^ and 10^5^, the flux is small, and infections do not start. When *S* = 10^6^ and 10^7^, both of the strains A and B have more than 100 bacteria of the *crossing* phenotype in the blood even if none of the cells switches, resulting in a mixed infection according to our definitions. At *S* = 10^7^, switching starts happening often, resulting in a faster activation of the immune response and clearance of the non-switched strain. Finally, when *S* = 10^8^, both strains are in large enough numbers to switch early on and at about the same time, resulting in mixed infections again, but now in the *growing* phenotypes.

**Figure 3.**
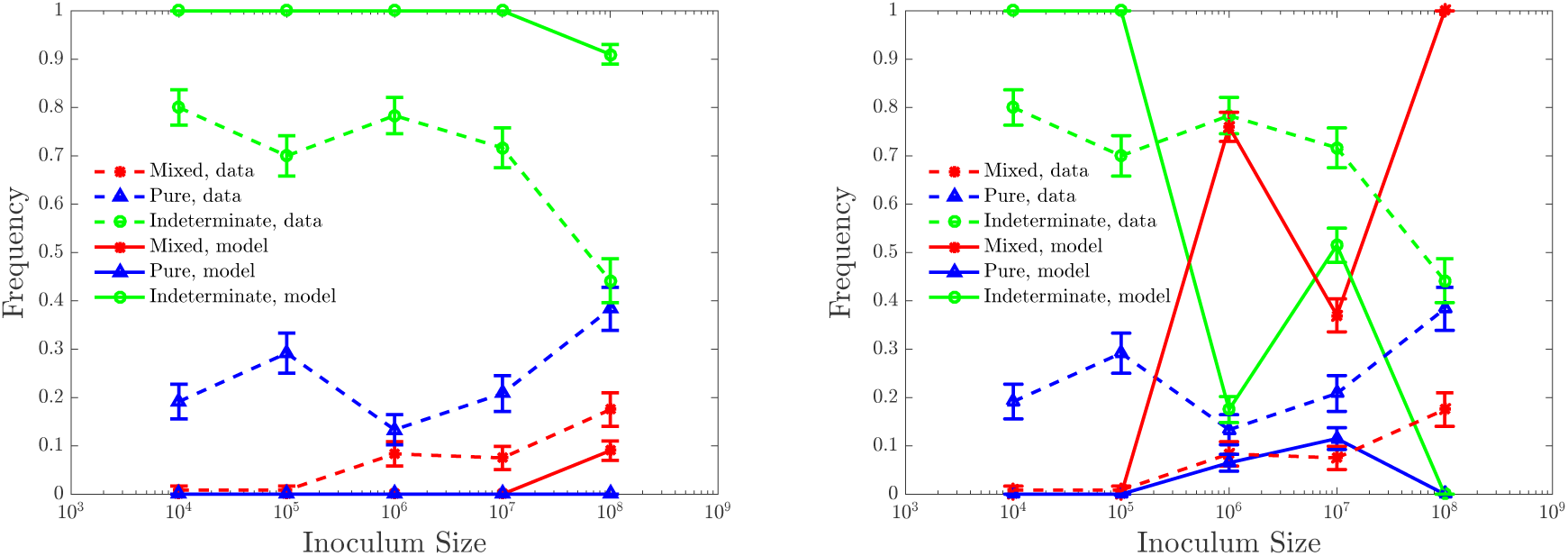
Maximum likelihood fits of the colloquial model. We show two different local maxima in the parameter space with nearly equivalent likelihoods. Neither set of parameters provides good fits. In this and subsequent figures, error bars on model predictions are given by standard deviations of results from 200 simulations. For the parameter values in the left panel (*ρ* ≈ 1.7·10^−5^h^−1^, *g*_*G*_ = 1.0 h^−1^, *r* ≈ 6.1·10^−6^h^−1^ cells^−1^, *γ* ≈ 240 h^−1^, *N*0 *≈* 1.8 · 10^9^ cells, *α*_*j*_ ≈ 4.5 · 10^−5^ h^−1^, *d* ≈ 7.2 · 10^−4^ h^−1^, *d*_*I*_ ≈ 1.7 · 10^−6^ h^−1^), infections do not establish until very large inoculums. For the right panel (*ρ* ≈ 1.7 · 10^−5^ h^−1^, *g*_*G*_ = 1.0 h^−1^, *r* ≈ 4.1 · 10^−8^ h^−1^ cells^−1^, *γ* ≈ 30.5 h^−1^, *N*0 ≈ 1.2 · 10^8^ cells, *α*_*j*_ ≈ 1.2 · 10^−5^ h^−1^, *d* = 0.01 h^−1^, *d*_*I*_ = 0.018 h^−1^), the need to establish pure infections over the four orders of magnitude in the inoculum size leads to a large number of mixed infections as well.

One can modify the model to make it fit better. Since rats have mucosal immunity [39, 40, 41, 42, 43], one can hope that bacteria in the nasopharynx will be eventually cleared as well. The simplest way of modeling this is to say that the flux from the nasopharynx into the bloodstream has a finite duration *t*_*j*_, which itself is an unknown variable that needs to be fitted. Further, we notice that the natural (not caused by the immune system) death rate of bacterial cells and the death rate of immune cells in the fits in Fig. 3 are very small. Hence, to not simplify fitting with the introduction of the new time parameter, we set both of these parameters to zero (we verified that the fits do not improve dramatically when this condition is relaxed). The optimized fit for this model is shown in Fig. 4. The fit is clearly better than in Fig. 3, but it is still poor: to have pure infections at medium/high inoculums, the independent action hypothesis still requires no (or indeterminate) infections at small inoculums.

**Figure 4.**
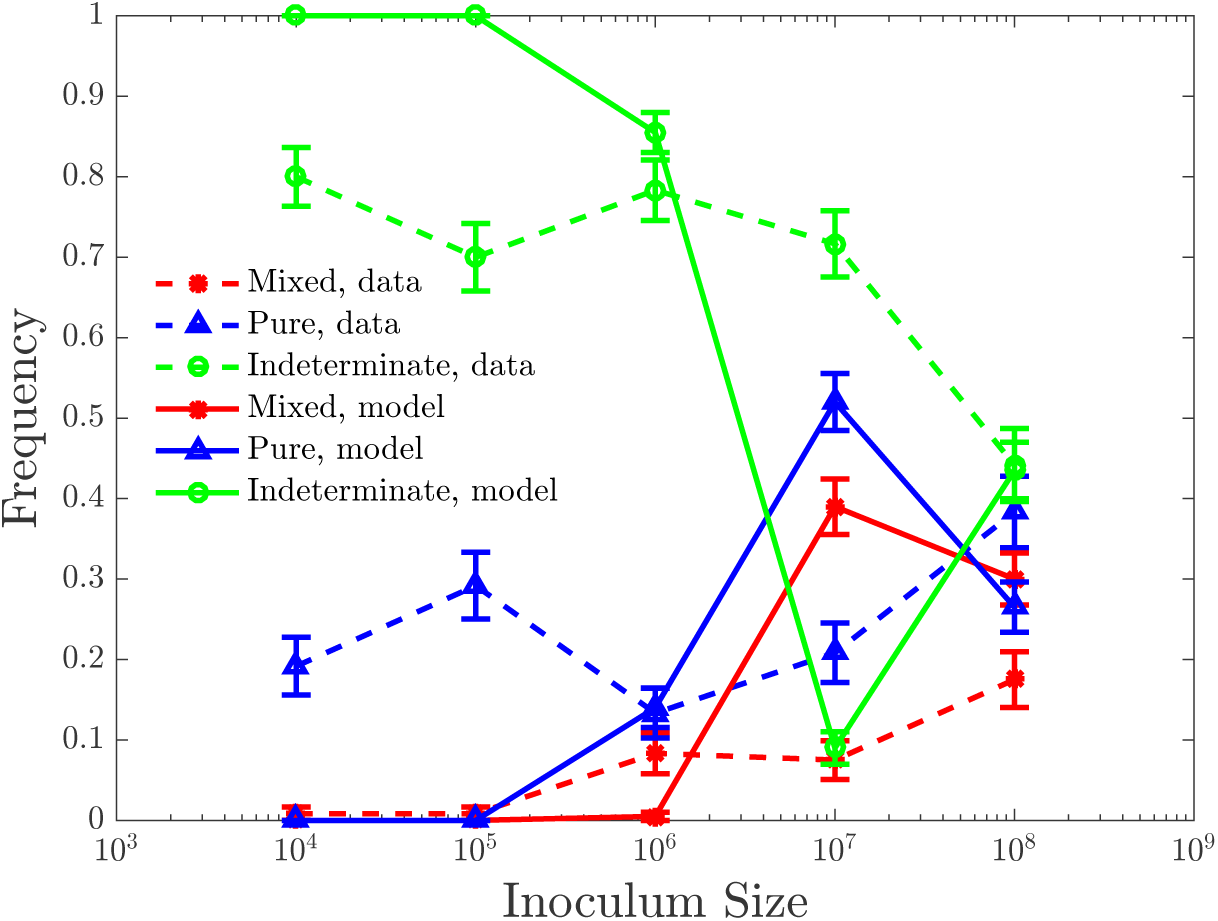
Maximum likelihood fits of the colloquial model with the limited time duration of the bacterial flux from the nasal cavity to the bloodstream. Natural bacterial cell death rate and immune cell death rate are set to zero. The duration of the flux is a fixed value for all inoculum sizes, *t*_*j*_ ≈ 4.1 h. Other optimized parameters are: *ρ* ≈ 3.5 · 10^−5^ h^−1^, *g*_*G*_ ≈ 0.96 h^−1^, *γ* ≈ 2.0 h^−1^, *r* ≈ 2.1 · 10^−7^ h^−1^ cells^−1^, *N*0 ≈ 1.0 · 10^6^ cells, *α*_j_ ≈ 1.4 × 10^−5^ h^−1^.

The key problem of the colloquial model is the experimentally observed weak dependence of the fractions of various infection types on the inoculum size. In other words, in these simple models, independent action means that the number of cells that attempt C→G switching in the blood scales with *S*. Thus the time to the first such switch would scale as 1/*S*. If the switch happens in the bulk of the 54 h experiment duration, an infection is established. Thus it is very hard to devise an independent action model that would have a non-negligible number of switches over 54 hrs at small inoculums, *S* = 10^4^, and yet would not have switches happening 100% of the time at large inoculums, *S* = 10^8^. The model must be modified so that the mean time to the first switch decreases slower than 1/*S*.

Interestingly, there is a straightforward biologically realistic modification of the model that achieves this. In many cases, the process of bacterial phenotypic switching is not determined by a one-step chemical reaction, but proceeds through a series of roughly equally slow steps. For example, the switching of *E. coli* to express PAP genes and become virulent [44, 45] can be modeled as a four-step reaction [33]. Such *n*-step activation ensures that the probability distribution of time to the complete switch in an individual bacterium goes as *∝ t*^*n*−1^ for small *t* [46, 47]. Then for *∝ S* bacteria, the expected time till the first of them switches is governed by the Weibull distribution, resulting in *St*^*n*−1^ *∝* 1, and *t ∝* 1/*S*^1/(*n*−1)^ [47]. In other words, the time to the first bacterium in a large population switching scales sublinearly with the inverse population size, offering a potential opportunity to explain the weak dependence on the inoculum size.

We implement and optimized this model in numerical simulations by introducing a series of phenotypic transitions C→G_1_ → … →G_*n*_, where only the last state, G_*n*_, grows fast, and the rest of the states share the growth/death rates with C. Random switching between the subsequent states was again governed by the Poisson dynamics, as in Eq. (4). We explored *n* = 2, 3, 4. Fig. 5 shows results of the optimization, where *n* = 3 (empirically best choice), and all transition rates in the chain C→G_1_ →G_2_ →G_3_ were the same (which results in the most sublinear dependence of the switch time on *S*). Further, since in this model switching takes extended time, the first bacteria to cross over to blood from the nasopharynx will be the ones switching, and hence it makes little difference for the switching statistics if the flux has a limited duration. At the same time, stopping of the flux into the blood results in a lower concentration of the non-switched strain, making it easier to develop pure infections at 54 hrs. Therefore, we inherit the value *t*_*j*_ ≈ 4.1 hrs from the 1-step model. Clearly, the quality of fit improves dramatically compared to the 1-step model, and yet the fits are still far from perfect: mixed and pure infections go hand-in-hand, and to have no mixed infections at *S* = 10^4^ requires having no infections at all at this inoculum. This illustrates a fundamental problem of the multi-step switching mechanism: while the time to a switch, indeed, scales sublinearly with 1/*S*, the standard deviation of this time falls off very quickly, making the switching nearly deterministic [47]. Thus both strains switch at about the same time, and typically either both develop into an infection (mixed outcome) or none does (indeterminate outcome).

**Figure 5.**
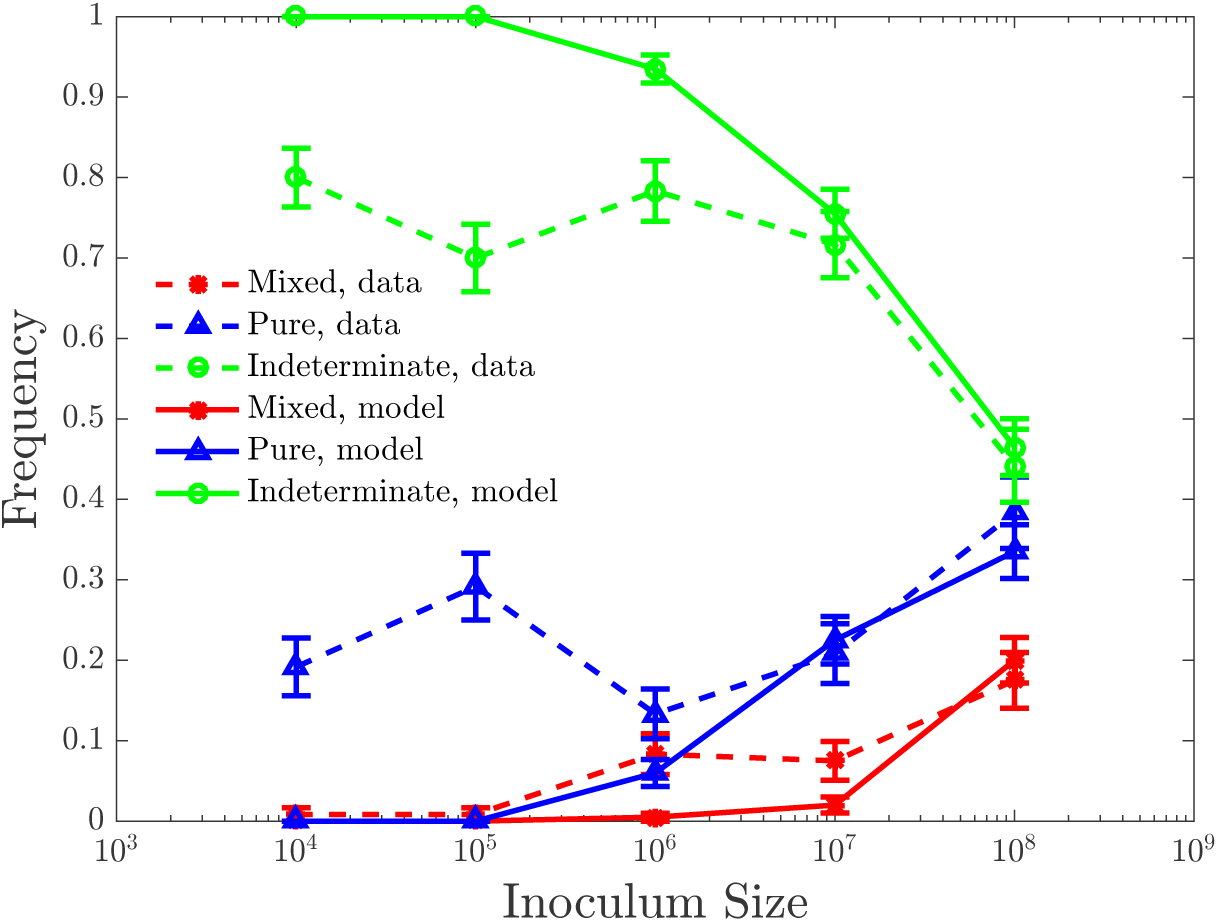
Maximum likelihood fits for the colloquial model with the limited bacterial flux duration and three-step switching. Natural bacterial cell death rate and immune cell death rate are set to zero. The duration of the flux is a fixed value for all inoculum sizes, *t*_*j*_ ≈ 4.1 h. Optimized parameter values are *ρ* ≈ 0.0014 h^−1^, *g*_*G*_ ≈ 1.1 h^−1^, *γ* ≈ 9.7 h^−1^, *r* ≈ 1.1 · 10^−7^ h^−1^ cells^−1^, *N*_0_ ≈ 1.0 · 10^9^ cells, *α*_*j*_ ≈ 3.4 · 10^−4^ h^−1^.

In summary, the independent action model, even augmented by multi-step switching and finite bacterial flux duration, seems to be incapable of explaining the observed experimental data.

### 3.2. Beyond the independent action model

The independent action hypothesis is implemented in our model, in part, by an assumption that the bacterial flux from the nasopharynx to blood is proportional to the inoculum size, *S*. We consider multiple extensions of the colloquial model that break this assumption.

First, we tried the model where the flux is independent of *S*, *j* = *α*_*j*_, which must be optimized. However, then the flux duration scales nonlinearly with *S*, *t*_*j*_ = *α*_*t*_S^*b*_*t*_^. The logic behind this model is that there might be purely physical constraints on how many bacteria can cross the tissues between the two body compartment per unit time, and this bandwidth can be saturated even at moderate inoculums. At the same time, it could take the mucosal immunity a longer time to clear a larger inoculum. We retain the 1-step switching model, because the higher variability of the switching time within this model allows for easier establishment of pure infections. The maximum likelihood results are shown in Fig. 6. While imperfect, the fits are surprisingly good, able to sustain pure, mixed, and indeterminate infections over the entire range of *S*. However, *b*_*t*_ ≈ 0.1 is very small, so that the duration of the bacterial flux is maximally ∼ 3 hours, which makes it hard to imagine physiological mechanisms that would create it.

**Figure 6.**
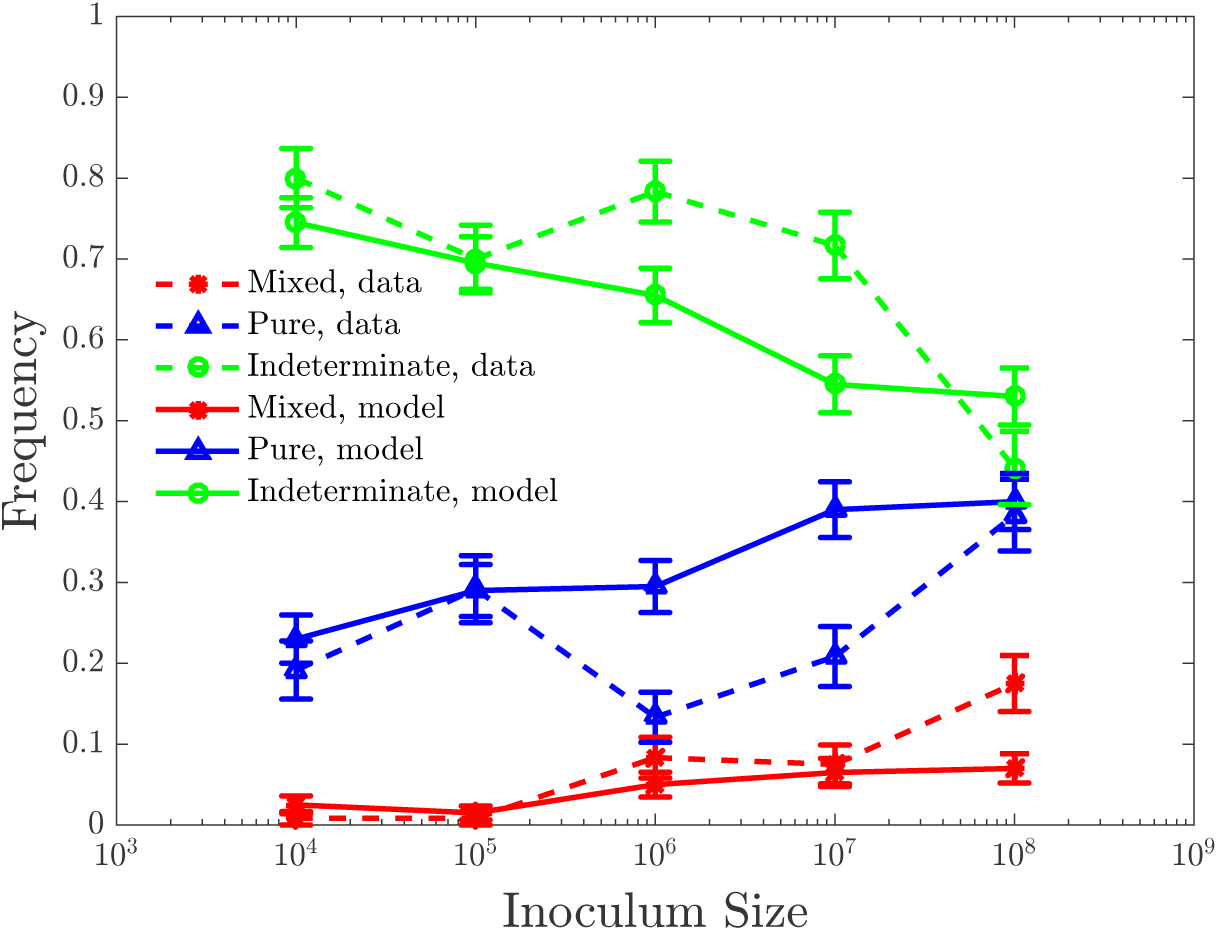
Maximum likelihood fits for the non-independent action model with the *S*-dependent flux duration. The fitted model has *j* = *α*_*j*_ = const, and *t*_*j*_ = *α*_*t*_*S*^*b*_*t*_^. This model provides much better fits than all of the variants of the independent action model we have tested. The optimized parameters are *ρ* ≈ 6.1 · 10^−6^ h^−1^, *g*_*G*_ ≈ 0.55 h^−1^, *N*0 ≈ 1.0·10^7^ cells, *γ* ≈ 2.0 h^−1^, *r* ≈ 2.3·10^−6^ h^−1^ cells^−1^, *α*_*t*_ ≈ 0.5 h^−1^, *b*_*t*_ ≈ 0.1, *α*_*j*_ ≈ 3.3 · 10^3^ h^−1^.

Another way to model non-independent action is the model where the duration of the bacterial flux *t*_*j*_ is fixed and independent of *S*, but *j* = *α*_*j*_S^*b*_*j*_^. Optimized dynamics for this model is shown in Fig. 7, providing clearly the best fit to the experimental data of the ones we have seen so far. Interestingly, the fitted values of the parameters in this model are biologically realistic, resulting, for example, in bacterial fluxes of 10^2^ ∼ 10^3^ cells/h, and *t*_*j*_ ≈ 34 hrs, longer than 1 day.

**Figure 7.**
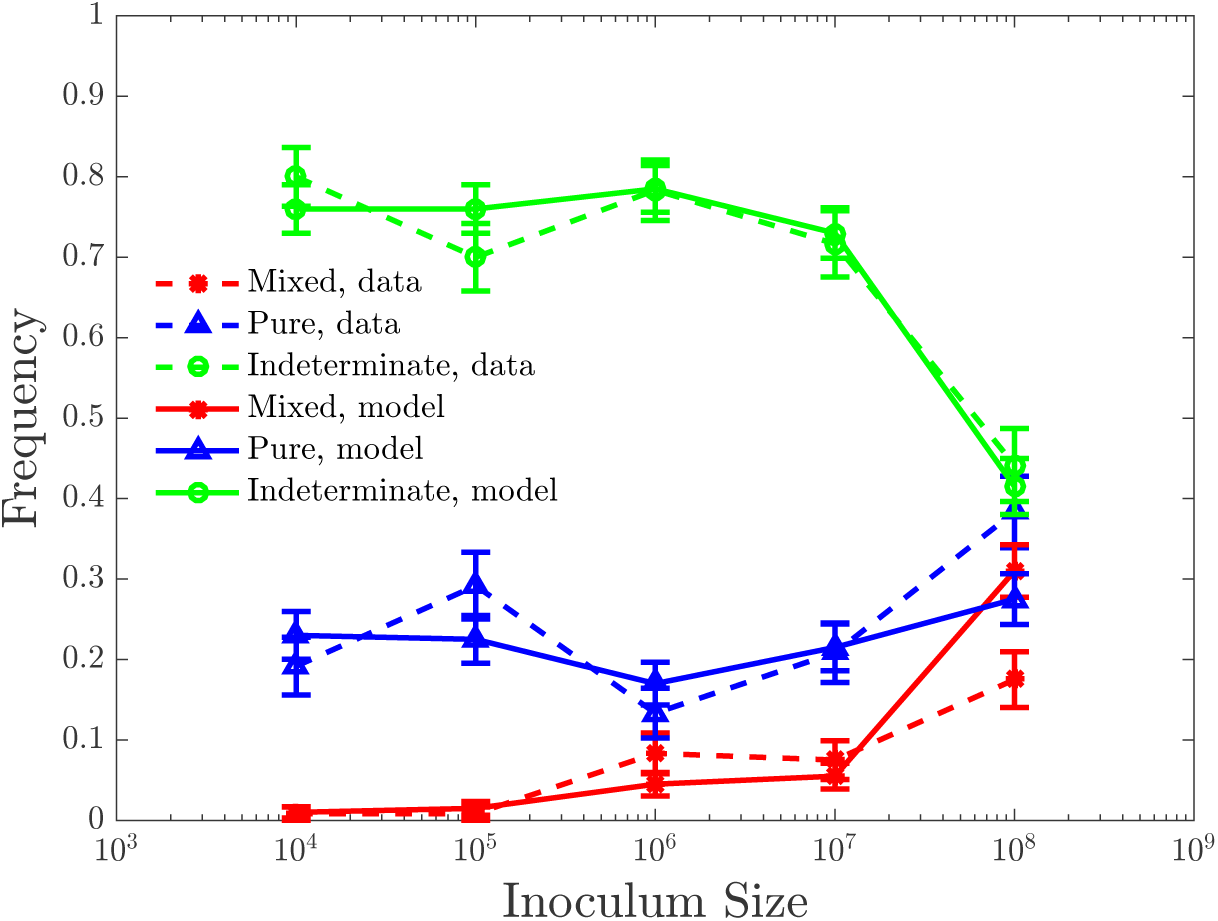
Maximum likelihood fits for the non-independent action model with the sublinear dependence of the magnitude of the bacterial flux on *S*. We use *j* = *α*_*j*_S^*b*_*j*_^ with a fixed *t*_*j*_. The optimized parameters are: *ρ* ≈ 5.7 · 10^−6^ h^−1^, *g*_*G*_ ≈ 1.0 h^−1^, *γ* ≈ 3.2 h^−1^, *r* ≈ 3.1 · 10^−6^ h^−1^ cells^−1^, *N*0 ≈ 1.0 · 10^6^ cells, *α*_*j*_ ≈ 7 h^−1^, *b*_*j*_ ≈ 0.37, and finally *t*_*j*_ ≈ 34.0 h, which is essentially equivalent to saying that the bacterial flux is temporally unconstrained.

## 4. Discussion

In this investigation, we built mathematical models of the early *Hib* stages of infections started by a culture with two equally invasive strains. The models have to account for the following broad observations: (1) both strains are present in the blood soon after inoculation into nasal passages; (2) both strains are either cleared from the blood within 54 hrs post inoculation, or are equally likely to be responsible for monoclonal bacteremias; and (3) the weak dependence of the likelihood of invasive infections on the inoculum size, i. e., a few-fold difference in the likelihood of bacteremia for a 10^4^-fold change in the size of the inoculum. The colloquial hypothesis tries to explain these effects as variations on the theme of independent action of bacteria in establishing infections. In such explanations, one commonly augments the independent action by stochasticity of phenotypic transitions and by interactions with the immune system. In this interpretation, both bacterial clones inoculated into the nasal passages cross into the bloodstream, at which point one individual randomly switches into a more invasive phenotype and activates the immune system. The immune system then clears the non-switched strain, resulting in monoclonal infections.

We analyzed this colloquial hypothesis quantitatively, starting with its simplest realization. Further, we considered additional variants that involved more complex (and hence statistically different) switching between the crossing and the growing phenotypes, or effects of mucosal immunity, which can clear the nasopharyngeal infection and stop bacterial flux into the bloodstream a few hours into the experiment. None of these modifications proved sufficient to explain the experimental data, and, in particular, to account for the weak dependence of the likelihood of the infections on the inoculum size. In contrast, when we forwent the independent action assumption and allowed the flux of bacteria into the bloodstream to depend sublinearly on the inoculum size, the fits to the data became very good. Thus our analysis suggests that the hypothesis of independent action may be violated in the case of early establishment of bacterial infections. Note that classic investigations of independent action [4] tested the hypothesis against *synergistic* effects. Here we argue that the non-independent action effects are redundant—the probability of a single bacterium to establish an infection decreases with the inoculum size. This is surprising, and certainly goes against ideas in the quorum sensing literature [48], where an infection is established synergistically when the number of bacteria crosses a certain minium density threshold.

Our best model suggests that the flux of bacteria from the nasopharyngeal inoculation to the bloodstream scales as the inoculum size to the ∼ 0.37 power. The data that we have been able to find in the literature is not sufficient to provide an empirical basis for the mechanism of this scaling: the physical structure of the animal tissues and bacterial colonies, the fluid dynamics of the bacterial culture in the nasopharyngeal cavity, interactions of bacteria with the immune system, or interactions of bacteria among themselves could all play a role. The closeness of the exponent to 1/3 is also interesting, suggesting that maybe a certain modification of the 3-step stochastic switching model, similar to that studied in Fig. 5, could play a role as well.

While informative at some level, negative results alone rarely appear in publications nowadays [49]. Beyond obvious social pressures, there are functional reasons for this as well: one can never be sure that the negative result is meaningful, rather than due to not trying hard to find a positive agreement between a hypothesis and data. For example, in our study, we cannot be sure that we have explored the parameter space well enough, and that we have tried all simple, reasonable modifications to the original colloquial model to argue that the independent action theory cannot explain the data. We can only say with certainty that *we have not been able to reconcile* the independent action theory with the data. It will require additional experimental and theoretical investigations to understand if and under which conditions the independent action hypothesis is, indeed, violated in early infections. The model proposed here would suggest that the bacterial concentration in the blood soon after inoculation should scale sublinearly with the incoculum size, which is relatively straight forward to check experimentally. Further, repeating the experiments in immune compromised rats, with no innate immunity, should result in no pure infections even at small inoculums. Moreover, due to the sublinear dependence, the probability of transmission of the disease among individuals should depend weakly on the strength of the infection in an infected host, which is bound to have public health implications, and is also experimentally testable. We hope that our study will spur such future investigations, in *Hib* and in other infections.

## Acknowledgements

This work was partially supported by the James S. McDonnell Foundation grant No. 220020321 (IN), by the National Science Foundation grant No. PoLS-1410978 (IN), and by National Institutes of Health grant No. GM098175 (BL). We thank Rustom Antia and Richard Moxon for illuminating discussions.

